## Supplemental material for "Effects of parental care on skin microbial community composition in poison frogs"

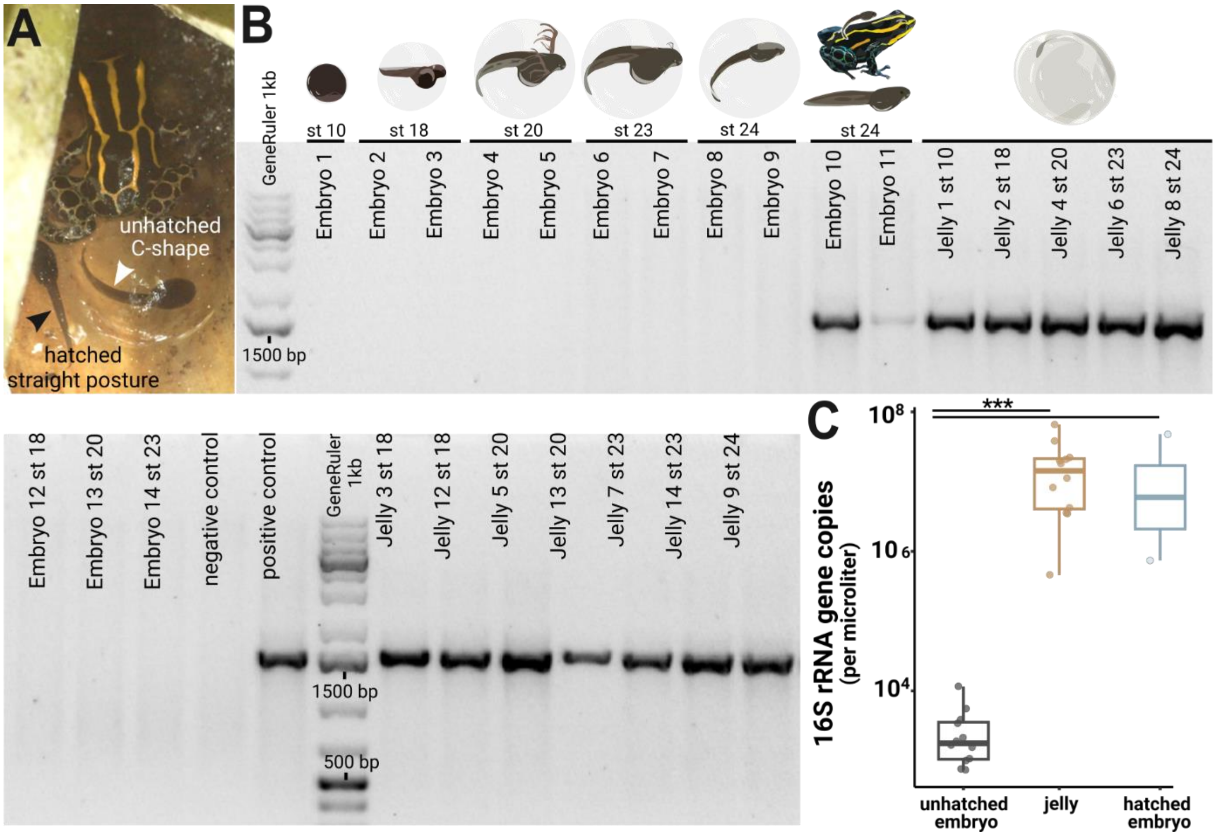


**Supplementary Figure 1: Microbes colonize a poison frog embryo after hatching from the vitelline membrane**. **(A)** Two tadpoles on the day of transport by their caregiver. Before hatching, tadpoles' posture is restricted by the transparent vitelline membrane, and they adopt a characteristic C-shape (right tadpole). After tadpoles hatch from the vitelline envelope, they display a straightened posture (left tadpole). Picture credit: Daniel Shaykevich. **(B)** DNA was isolated from whole embryos and jellies of different developmental stages. The 16S rRNA gene was amplified with a broad range PCR using primer specific for the full 16S rRNA region (27F and 1492R, ~1500bp). Amplification products were visualized on a 2% agarose gel. **(C)** 16SrRNA gene copy numbers were quantified in jellies, hatched and unhatched embryos using a digital droplet PCR (ddPCR). Variations in copy numbers between the three groups were analyzed with a Kruskal-Wallis test with Benjamini-Hochberg correction followed by a Dunn test. Significances <0.001 are indicated by ***.

**Supplementary Table 1: Copy number variations of the 16S rRNA gene between embryos and jelly detected by ddPCR.** DNA extracted from jellies, hatched and unhatched embryos was analyzed using a QX200 AutoDG Droplet Digital PCR system (Bio-Rad) with universal 16S rRNA primers (331F/797R) and 16S rRNA FAM probes. Displayed copy numbers were corrected for dilution factor, extraction elution volume, and sample ddPCR volume and represent 16S rRNA copy numbers/µl present in the jelly or embryo of one *Rv* egg. Mean copy numbers/µl and standard deviations were calculated for each Gosner stage.

| ID | group | copies/µl | mean copies/µl/stage ± STD | Gosner stage |
|---|---|---|---|---|
| Embryo 1 | unhatched | 3463 | 3463 (~3x$10^3$) | st 10 |
| Embryo 2 | unhatched | 2161 | | st 18 |
| Embryo 3 | unhatched | 999 | 1611 (~1 x $10^3$) ± 583 | st 18 |
| Embryo 12 | unhatched | 1673 | | st 18 |
| Embryo 4 | unhatched | 3853 | | st 20 |
| Embryo 5 | unhatched | 766 | 1896 (~2 x $10^3$) ± 1702 | st 20 |
| Embryo 13 | unhatched | 1068 | | st 20 |
| Embryo 6 | unhatched | 1886 | | st 23 |
| Embryo 7 | unhatched | 742 | 4837 (~4 x $10^3$) ± 5957 | st 23 |
| Embryo 14 | unhatched | 11584 | | st 23 |
| Embryo 8 | unhatched | 5561 | 3554 (~4 x $10^3$) ± 2837 | st 24 |
| Embryo 9 | unhatched | 1548 | | st 24 |
| Embryo 10 | hatched | 48159335 | 24448337 (~2 x $10^7$) ± 33532414 | st 24 |
| Embryo 11 | hatched | 737340 | | st 24 |
| Jelly 1 | jelly | 37998434 | 18029604 (~1 x $10^7$) ± 583 | st 10 |
| Jelly 2 | jelly | 3550824 | | st 18 |
| Jelly 3 | jelly | 4235222 | | st 18 |
| Jelly 12 | jelly | 21107921 | | st 18 |
| Jelly 4 | jelly | 21990138 | | st 20 |
| Jelly 5 | jelly | 457417 | | st 20 |
| Jelly 13 | jelly | 19812023 | | st 20 |
| Jelly 6 | jelly | 8318251 | | st 23 |
| Jelly 7 | jelly | 18013704 | | st 23 |
| Jelly 14 | jelly | 11299376 | | st 23 |
| Jelly 8 | jelly | 66115451 | | st 24 |
| Jelly 9 | jelly | 3456484 | | st 24 |

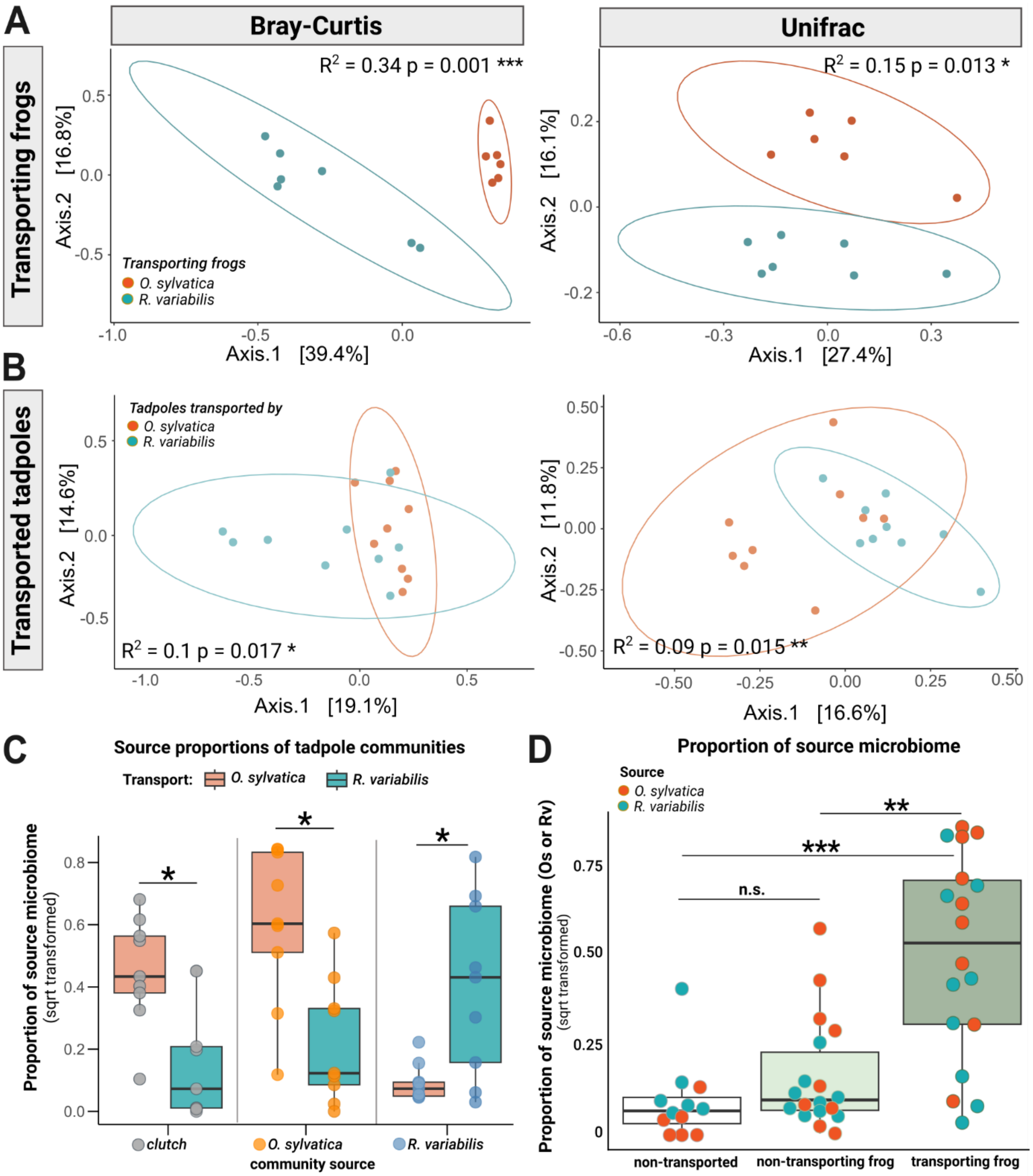


**Supplementary Figure 2: Transporting frogs serve as source of microbes for transported tadpoles**. **(A)** Communities of frog species *Rv* (blue) and *Os* (orange) that transported tadpoles cluster distinctly from each other in a principal coordinate analysis of Bray Curtis (left panel) and Unifrac (right panel) distances. **(B)** After 6 hours of transport, communities of tadpoles transported by *Rv* and *Os* overlap but cluster distinctly. **(C)** In a separate analysis, SourceTracker was trained on the communities of non-transported siblings as an additional source to assess whether microbes associated with transported tadpoles had been acquired from the clutch; this approach also served to test the consistency of our findings across different source configurations. Source proportions (clutch: grey dots, *Os*: orange dots, *Rv*: blue dots) were determined for tadpoles transported by *Os* (orange boxes) or *Rv* (blue boxes) and compared with a pairwise Wilcoxon test. P-values were adjusted using Benjamini Hochberg correction. All source proportions were square root transformed for plotting. **(D)** We used Sourcetracker to identify the sources of taxa (identified with 16S v4 specific amplicon sequences) that had been acquired by tadpoles. The function was trained on communities of adult *Rv* and *Os* that had served as caregivers. Source proportions of both species (*Os*: orange dots and *Rv*: blue dots) were determined for each tadpole (N = 24), resulting in 2 data points per tadpole. Proportions were then grouped to display either (1) proportions of the transporting species in transported tadpoles (*Rv* proportions in tadpoles transported by *Rv* and *Os* proportions in tadpoles transported by *Os*) (dark green), or (2) proportions of the non-transporting species on transported tadpoles (indicating *Rv* proportions in tadpoles transported by *Os* and *Os* proportions in tadpoles transported by *Rv*) (light green), or (3) proportions of both species in non-transported tadpoles (indicating *Rv* and *Os* proportions in non-transported tadpoles) (white). Proportions were compared with a Kruskal-Wallis test with Benjamini-Hochberg correction. Source proportions were square root transformed for plotting.

**Supplementary Table 2: Source proportions of *Os* and *Rv* communities in the microbiome of tadpoles transported by *Os*, and their sibling transported by *Rv* or non-transported.** Siblings belonging to the same clutch are indicated by color blocks. Abbreviations: exp group = experimental group, syl = transported by Os, var = transported by Rv, control = non- ransported, transp = transported

| Tadpole ID | Exp group | Family ID | % Os | % Rv | % more Os than in Rv-transp sibling | % more Os than in non-transp sibling |
|---|---|---|---|---|---|---|
| 1 | control | 2 | 1.759 | 16.434 | | |
| 2 | syl | 2 | 41.163 | 6.570 | 39.33 | 39.44 |
| 3 | syl | 2 | 70.311 | 1.088 | 68.478 | 68.552 |
| 4 | var | 2 | 1.833 | 2.640 | | |
| 5 | control | 3 | 0.000 | 2.132 | | |
| 6 | syl | 3 | 72.973 | 1.351 | 4.27 | 72.973 |
| 7 | var | 3 | 32.766 | 18.816 | | |
| 8 | control | 4 | 0.175 | 0.526 | | |
| 9 | syl | 4 | 68.349 | 0.272 | 57.965 | 68.174 |
| | | | | | 59.978 | 9.7 |
| 10 | var | 4 | 8.371 | 17.400 | | |
| 11 | var | 4 | 10.385 | 68.956 | | |
| 12 | control | 5 | 0.000 | 0.900 | | |
| 13 | syl | 5 | 0.874 | 0.291 | 0.872 | 0.874 |
| 14 | syl | 5 | 34.730 | 0.811 | 34.728 | 34.73 |
| 15 | var | 5 | 0.002 | 9.623 | | |
| 16 | control | 6 | 0.250 | 0.375 | | |
| 17 | syl | 6 | 50.586 | 0.554 | 49.876 | 5.586 |
| 18 | var | 6 | 0.710 | 0.118 | | |
| 19 | control | 7 | 0.000 | 0.678 | | |
| 20 | syl | 7 | 22.593 | 2.222 | 22.534 | 22.593 |
| 21 | syl | 7 | 9.381 | 0.442 | 7.158 | 9.381 |
| 22 | var | 7 | 0.059 | 43.985 | | |

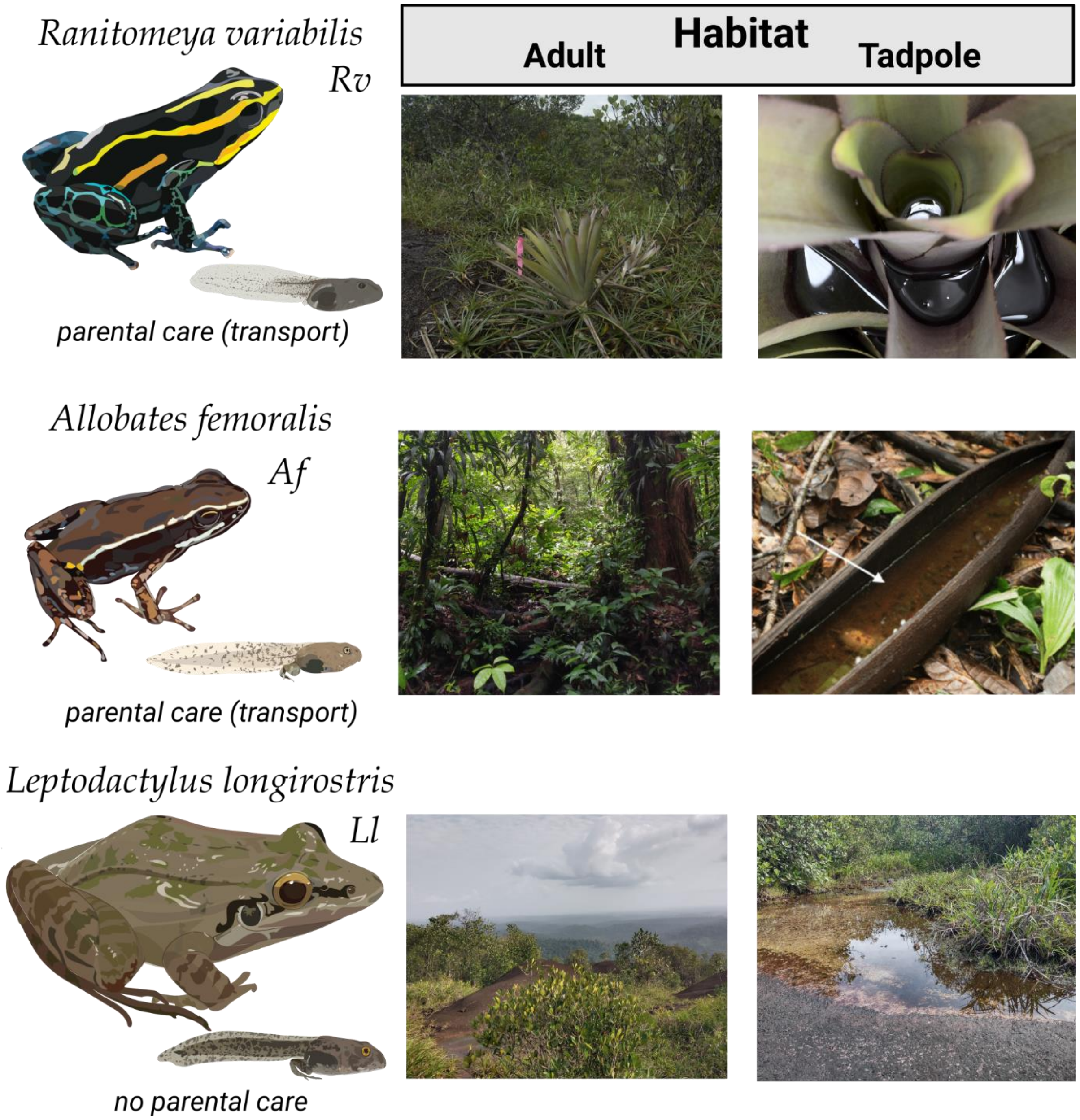


**Supplementary Figure 3: Study species and their habitat.** We compared the skin microbiome of three anuran species: *Rv* (upper panel) and *Af* (middle panel) are poison frogs that transport their offspring but prefer different habitat types (leaf litter in primary forest vs. rock savanna on a mountain plateau). While both transport their offspring, *Af* tadpoles grow up in groups and *Rv* tadpoles are cannibalistic and grow up in individual leaf axil pools. *L. longirostris* (lower panel) is a leptodactylid frog that cohabits the rock savanna with *Rv* but deposits its eggs in water without transporting its tadpoles. Picture of *Af* tadpole habitat published by Rojas & Pašukonis 2019.

**Supplementary Table 3: Taxonomic composition of microbiome samples from wild tadpoles and adults.** Total number of phyla and families encountered in each group were calculated from unrarefied data, averaged group values and standard deviations were calculated from a rarefied dataset. *Batrachochytrium dendrobatidis* (Bd) positive samples were extracted from an ITS sequencing dataset and confirmed with a nested PCR targeting Bd. Abbreviations: #: number, Bd +: number of samples positive for *Batrachochytrium dendrobatidis*, avg: average, inhibit.: inhibiting. Maximas are displayed bold, lowest number of *Bd* inhibiting taxa are marked grey.

| Species | Sample | N | Total # Phyla | Avg # Phyla | Total # Families | Avg # Families | Bd + | Avg # Bd inhibit. taxa |
|---|---|---|---|---|---|---|---|---|
| | A | 44 | 32 | 7.3 ± 3.9 | 265 | 26.4 ± 19.1 | 2 (4.5 %) | 5.7 ± 3.6 |
| ***R. variabilis*** | T | 21 | 25 | 9.1 ± 4.1 | 185 | 29.1 ± 16.2 | 1 (4.7 %) | 1.1 ± 0.5 |
| | W | 21 | **32** | **17.5** ± 3.2 | **283** | **73.7 ± 14.6** | - | 1.6 ± 1.6 |
| | A | 10 | 28 | 9.2 ± 3.5 | 208 | 34.7 ± 19.8 | 1 (10%) | 7.6 ± 6.0 |
| ***A. femoralis*** | T | 8 | 21 | 7.3 ± 2.5 | 104 | 29 ± 13.6 | 0 | 3.1 ± 2.0 |
| | W | 3 | **33** | **14.3** ± 1.5 | **267** | **75.3 ± 2.1** | - | 11.7 ± 4.0 |
| | A | 10 | 16 | 9.7 ± 2.9 | 112 | 31.9 ± 11.6 | 0 | 6.3 ± 4.4 |
| ***L. longirostris*** | T | 14 | 19 | 4.3 ± 2.0 | 98 | 11.6 ± 6.0 | 0 | 0.4 ± 0.5 |
| | W | 6 | **25** | **13.8** ± 1.3 | **172** | **66.5 ± 5.4** | - | 3.2 ± 1.2 |

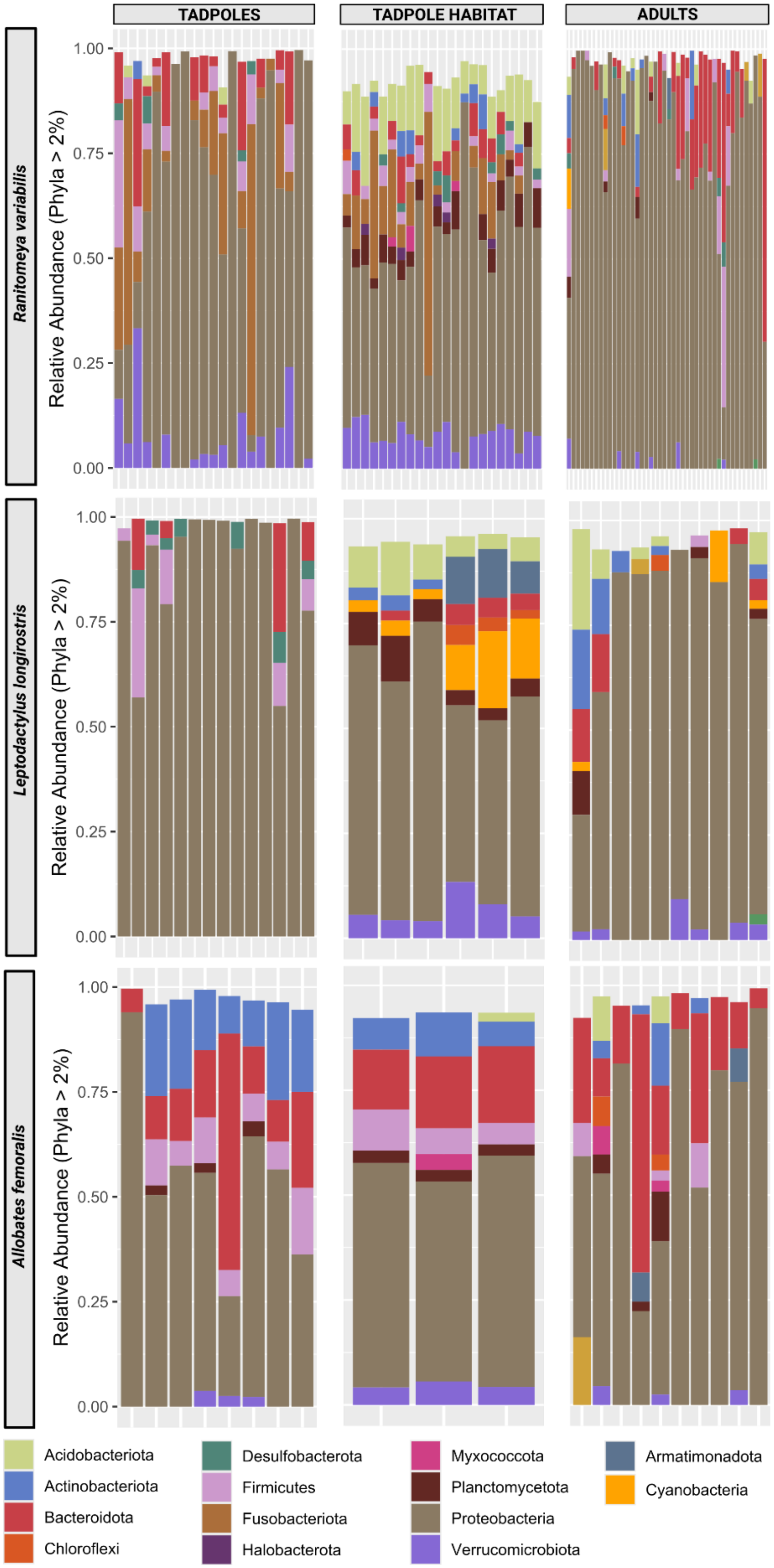


**Supplementary Figure 4: Phyla distribution per species and group.** Bars represent individual tadpoles, frogs or samples of the aquatic environment. We sampled triplicates of aquatic environments of group-living tadpoles occurring in the same pool. Colors represent phyla that were detected with a relative abundance of more than 2%.

**Supplementary Table 4: Statistical analysis of alpha diversity measures across species and life stages and environments.** Statistical tests for alpha diversity were performed on rarefied datasets. Significant ANOVAs were followed by Tukey post-hoc tests. Sample sizes for each comparison are provided under the respective group. Significance levels of adjusted p values are indicated by * (p < 0.05), ** (p < 0.01) or *** (p < 0.001). Abbreviations: T: tadpole, A: adult, *Rv*: *Ranitomeya variabilis*, *Af*: *Allobates femoralis*, *Ll*: *Leptodactylus longirostris*.

| Species | Comparison | Diversity measure | Test type | Df | Test Statistic | p.adj |
|---|---|---|---|---|---|---|
| *Rv* | A - T | Observed ASV | KW | 1 | $Chi^2$ = 4.7004 | 0.0905 |
| | (44 – 21) | Shannon | ANOVA | 1 | F = 0.5267 | 1.4121 |
| | | evenness | ANOVA | 1 | F = 0.0099 | 2.7636 |
| *Af* | A -T | Observed ASV | KW | 1 | $Chi^2$ = 0.19737 | 1.9707 |
| | (10 – 8) | Shannon | ANOVA | 1 | F = 0.1843 | 2.0202 |
| | | evenness | KW | 1 | $Chi^2$ = 0.95526 | 0.9852 |
| *Ll* | A – T | Observed ASV | KW | 1 | $Chi^2$ = 7.2612 | 0.0211* |
| | (10 – 10) | Shannon | ANOVA | 1 | F = 31.143 | < 0.001*** |
| | | evenness | ANOVA | 1 | F = 33.492 | < 0.001*** |
| *Rv – Ll* | T - T | Observed ASV | KW | 2 | $Chi^2$ = 10.158 | 0.0127* |
| | (21 – 10) | Shannon | ANOVA | 2 | F = 10.973 | 0.0020** |
| | | evenness | ANOVA | 2 | F = 10.982 | 0.0024** |
| *Af – Ll* | T - T | Observed ASV | KW | 2 | $Chi^2$ = 10.158 | 0.0149* |
| | (8 – 10) | Shannon | ANOVA | 2 | F = 10.973 | 0.0003** |
| | | evenness | ANOVA | 2 | F = 10.982 | 0.0003** |
| Rv - Ll | W – W | Observed ASV | ANOVA | 2 | F = 18.752 | 0.0037* |
| | (21 – 6) | Shannon | ANOVA | 2 | F = 11.205 | 0.0023* |
| | | evenness | ANOVA | 2 | F = 0.01104 | 0.0212* |
| Rv- Af | W – W | Observed ASV | ANOVA | 2 | F = 18.752 | < 0.001*** |
| | (21 – 3) | Shannon | ANOVA | 2 | F = 11.205 | 0.005* |
| | | evenness | ANOVA | 2 | F = 0.01104 | 0.1278 |
| Af - Ll | W – W | Observed ASV | ANOVA | 2 | F = 18.752 | < 0.001*** |
| | (3-6) | Shannon | ANOVA | 2 | F = 11.205 | 0.9936 |
| | | evenness | ANOVA | 2 | F = 0.01104 | 0.8436 |

**Supplementary Table 5: PERMANOVA for Principal Coordinate Analysis on Bray Curtis distances**. Adonis permutation was followed by a pairwise adonis to determine if communities of groups cluster distinctly. Significance levels of p and adjusted p values are indicated by * ($p < 0.05$), ** ($p < 0.01$) or *** ($p < 0.001$).

| Comparison | Df | SumsOfSqs | F Model | R2 | P value | Adj. p |
|---|---|---|---|---|---|---|
| *A_Rv* vs *A_Ll* | 1 | 1.992 | 7.796 | 0.130 | 0.001 *** | 0.036 * |
| *A_Rv* vs *A_Af* | 1 | 3.158 | 11.513 | 0.181 | 0.001 *** | 0.036 * |
| *A_Ll* vs *A_Af* | 1 | 2.081 | 7.941 | 0.306 | 0.001 *** | 0.036 * |
| *A_Rv* vs *T_Rv* | 1 | 6.659 | 27.721 | 0.306 | 0.001 *** | 0.036 * |
| *T_Ll* vs *A_Ll* | 1 | 4.089 | 29.143 | 0.570 | 0.001 *** | 0.036 * |
| *T_Af* vs *A_Af* | 1 | 2.338 | 9.503 | 0.373 | 0.001 *** | 0.036 * |
| *T_Ll* vs *T_Af* | 1 | 3.711 | 32.184 | 0.617 | 0.001 *** | 0.036 * |
| *T_Af* vs *T_Rv* | 1 | 3.671 | 20.619 | 0.433 | 0.001 *** | 0.036 * |
| *T_Ll* vs *T_Rv* | 1 | 5.779 | 38.646 | 0.539 | 0.001 *** | 0.036 * |
| *T_Af* vs *Af_water* | 1 | 0.842 | 6.322 | 0.413 | 0.013 * | 0.468 |
| *T_Rv* vs *Rv_water* | 1 | 4.704 | 26.064 | 0.389 | 0.001 *** | 0.036 * |
| *T_Ll* vs *Ll_water* | 1 | 3.221 | 32.362 | 0.643 | 0.001 *** | 0.036 * |

**Supplementary Table 6: Number of core taxa (genus level agglomerated) across different prevalence and abundance cutoffs.** Abbreviations: prev = prevalence, abd = abundance, aqu env = aquatic environment

| Sample | prev 100% (no abd) | prev 75%, abd 1% | prev 75%, abd 0.1% | prev 75% (no abd) |
|---|---|---|---|---|
| ***Rv* adults** | 0 | 1 | 1 | 2 |
| ***Rv* tadpoles** | 0 | 3 | 6 | 14 |
| ***Rv* aqu env** | 0 | 4 | 24 | 69 |
| ***Af* adults** | 0 | 0 | 4 | 7 |
| ***Af* tadpoles** | 0 | 7 | 7 | 7 |
| ***Af* aqu env** | 0 | 12 | 81 | 236 |
| ***Ll* adults** | 0 | 2 | 4 | 4 |
| ***Ll* tadpoles** | 0 | 2 | 2 | 6 |
| ***Ll* aqu env** | 0 | 6 | 46 | 110 |

**Supplementary Table 7: Identity of core genera across different occurrence and abundance cutoffs**. Prevalence of each genus in the aquatic environment of the respective tadpoles is indicated by "N" (no) or "Y" (yes). Colors indicate cutoff levels (yellow: prevalence > 75%, blue: prevalence > 75% and abundance > 0.1%, black: prevalence > 75% and abundance > 1%), $H_2O$ = aquatic environment.

| Sample | Core Genera | $H_2O$ |
|---|---|---|
| ***Rv* adults** | **"Rosenbergiella"** | **N** |
| | "Methylobacterium-Methylorubrum" | N |
| ***Rv* tadpoles** | **"Cetobacterium"** | **Y** |
| | **"Novosphingobium"** | **Y** |
| | **"Pelomonas"** | **Y** |
| | **"Bacteroides"** | **N** |
| | **"Alistipes"** | **N** |
| | **"Bacteria_Verrucomicrobiota_Verrucomicrobiae_Opitutales_Puniceicoccaceae "** | **N** |
| | "Rikenella" | N |
| | "Desulfovibrio" | Y |
| | "Clostridium_sensu_stricto_1" | Y |
| | "Tyzzerella" | Y |
| | "Monoglobus" | N |
| | "Ruminococcus" | N |
| | "Roseiarcus" | Y |
| | "Rhodomicrobium" | Y |
| ***Af* adult** | **"Candidatus_Hemobacterium"** | **N** |
| | **"Pedobacter"** | **N** |
| | **"Pseudomonas"** | **Y** |
| | **"Allorhizobium-Neorhizobium-Pararhizobium-Rhizobium"** | **N** |
| | "Bacteria_Planctomycetota_Planctomycetes_Gemmatales_Gemmataceae_uncultured" | Y |
| | "Bradyrhizobium" | Y |
| | "Burkholderia-Caballeronia-Paraburkholderia" | **Y** |
| ***Af* tadpoles** | **"Mycobacterium"** | **Y** |
| | **"Bacteria_Bacteroidota_Bacteroidia_Bacteroidales_Barnesiellaceae "** | **Y** |
| | **"Rikenella"** | **Y** |
| | **"Dinghuibacter"** | **Y** |
| | **"Dechloromonas"** | **Y** |
| | **"Denitratisoma"** | **Y** |
| | **"Aeromonas"** | Y |
| ***Ll* adults** | **"Methylobacterium-Methylorubrum"** | **Y** |
| | **"Rhodomicrobium"** | **Y** |
| | **"Burkholderia-Caballeronia-Paraburkholderia"** | **Y** |
| | **"Candidatus_Xiphinematobacter"** | **Y** |
| ***Ll* tadpoles** | **"Pandoraea"** | **Y** |
| | **"Curvibacter"** | **Y** |
| | "Desulfovibrio" | Y |
| | "Tyzzerella" | Y |
| | "Aquitalea" | Y |
| | "Rikenella" | N |

### Table S8: ANCOMBC Analysis -provided as Excel file

**Supplementary Table 9: Identity, relative contribution of the caregiver's microbiome, relative abundance in tadpole communities, and source-pool presence of ASVs shared between transported tadpoles and their adult caregivers.** Relative contribution of the caregiver's microbiome was determined with Sourcetracker. Abundances over 10% are indicated bold. Abbreviations: T = tadpole, A = adult, relA = relative abundance.

| tadpole ID | Proportion of caregiver microbiome (Sourcetracker) | ASVs shared T-A | Presence in source: adult (A), adult or water (A/W) | ASV | relA (%) | Presence in caregivers 10 most abundant ASV |
|---|---|---|---|---|---|---|
| **transported 1** | 1% | 2 | A<br>A/W | 102:Akkermansia,<br>144:Enterobacteriaceae | 1.847<br>1.450 | yes<br>no |
| **transported 2** | 17% | 1 | A/W | 144:Enterobacteriaceae | **19.258** | yes |
| **transported 3** | 0 % | 1 | A/W | 312:Elsteraceae | 0.125 | no |
| **transported 4** | 0 % | 0 | 0 | - | 0 | - |
| **transported 5** | 0 % | 0 | 0 | - | 0 | - |
| **transported 6** | 15 % | 2 | A<br>A/W | 138:Aquitalea,<br>421:Methylocella | 0.842<br>**18.050** | yes<br>yes |
| **transported 7** | 0% | 1 | A/W | 88:Methylobacterium-Methylorubrum | 0.020 | yes |
| **transported 8** | 9% | 1 | A | 911:Staphylococcus | **10.329** | yes |
| **transported 9** | 0% | 2 | A<br>A/W | 28:Akkermansia,<br>3092:Pseudomonas | 1.378<br>0.090 | no<br>no |
| **transported 10** | 0% | 1 | A/W | 219:Acinetobacter | **6.259** | yes |

**Supplementary Table 10: Scaled absolute abundance of ASVs shared between transported tadpoles and their adult caregivers as identified by Sourcetracker.** 16S rRNA copy numbers per microliter were determined by digital PCR (dPCR) using the QIAcuity system with the same primer set used for the v4 amplicon dataset and quantified in triplicate across different dilutions; reported values represent means with standard deviations. Relative abundances of shared ASVs, obtained from the v4 amplicon sequencing dataset, were used to scale total 16S rRNA gene copy numbers per microliter, yielding ASV-specific scaled absolute abundances. Scaled abundances corrected for the elution volume are given as copies per tadpole. Abbreviations: relA = relative abundance, absA = absolute abundance, STD = standard deviation.

| tadpole ID | ASV | relA (%) | scaled absA (copies/µl) | scaled absA (copies/tadpole) | mean 16S rRNA copies/µl ± STD |
|---|---|---|---|---|---|
| **transported 1** | 102:Akkermansia,<br>144:Enterobacteriaceae | 1.847<br>1.450 | 383<br>301 | 19141<br>15027 | 20727 ± 6069 |
| **transported 2** | 144:Enterobacteriaceae | 19.258 | 156 | 7804 | 810 ± 212 |
| **transported 6** | 138:Aquitalea,<br>421:Methylocella | 0.842<br>18.050 | 161<br>3447 | 8039<br>172326 | 19094 ± 458 |
| **transported 8** | 911:Staphylococcus | 10.329 | 728 | 36408 | 7050 ± 1251 |

**Supplementary Table 11: The presence of ASVs shared between transported tadpoles and their caregivers in non-transported tadpoles.**

| tadpole ID | Caregiver ASVs detected in tadpole | ASVs | ASVs present in water (out of 10 ASVs) |
|---|---|---|---|
| **non-transported 1** | 1 | 144:Enterobacteriaceae | 7 |
| **non-transported 2** | 0 | 0 | 6 |
| **non-transported 3** | 0 | 0 | 7 |
| **non-transported 4** | 1 | 144:Enterobacteriaceae | 6 |
| **non-transported 5** | 1 | 144:Enterobacteriaceae | 6 |
| **non-transported 6** | 0 | 0 | 6 |
| **non-transported 7** | 1 | 144:Enterobacteriaceae | 7 |
| **non-transported 8** | 0 | 0 | 6 |
